# An Inflammation Centered Perspective to the Mechanisms and Interactions Related to Vascular Cognitive Impairment

**DOI:** 10.1101/2024.09.19.613013

**Authors:** Melisa Ece Zeylan, Simge Senyuz, Ozlem Keskin, Attila Gursoy

## Abstract

A major health burden for the elderly, vascular cognitive impairment (VCI) is a disease that combines cognitive (CD) and cardiovascular (CVD) components. The molecular mechanisms underlying this disease are poorly understood, and our work attempts to bridge this knowledge gap by building protein-protein interaction (PPI) networks of CD and CVD. Our earlier research not only showed how well these two primary components work together, but also hinted at the potential role of inflammation in the development of VCI. For this reason, we decided to examine the relationship between inflammation and VCI in further detail.We identified the top three most connected clusters, which could represent significant modules, enriched these clusters with alternative conformations, and used PRISM to predict the interactions between the conformations. We proposed putative VCI-related interactions, such as NFKBIA-RELA and the proteasome complex, as well as their effects. The five interactions that we discovered have a higher predicted binding affinity when one of the conformations is mutated: LTF-SNCA, FGA-LTF, UBE2D1-VCP, ERBB4-INS, and NFE2L2-VCP. Additionally, since VCP has a conformational mutation linked to dementia, we proposed that the cancer-related protein BRCA1 may have implications for VCI. BRCA1’s interaction with both wild-type and mutant XRCC4 and LIG4 suggests the significance of the DNA damage response pathway which can be shared between VCI and cancer.Altogether, our results suggest various pathways and interactions that can act as targets for therapeutic interventions or early diagnosis of VCI.

## INTRODUCTION

Vascular cognitive impairment (VCI) is a severe problem that is linked to several cognitive deficiencies, such as executive function, memory and attention impairment [1]. VCI is a complex disease that is impacted by many factors, with the significant contribution of cardiovascular risk factors and cardiovascular disease subtypes (CVD) [2, 3, 4]. The severe implications of this disease make it crucial to detect biomarkers and pathways that can serve to understand the disease. We can study such complex diseases with a systems biology approach by constructing networks regarding their two main components, CVD and cognitive decline (CD) and analyzing them.

Furthermore, VCI is a condition heavily influenced by inflammation [5]. A number of adverse circumstances, such as infection, ischemia, and hypoxia, can cause inflammatory reactions. These reactions can lead to overactivated neuroinflammatory responses, causing apoptosis, blood-brain barrier damage (BBB) and other pathological conditions[6]. According to Exel et al., this inflammation is especially harmful to people who have CVD because of its association with CD[7]. Therefore, this undeniable effect of inflammation on VCI makes it crucial to take an inflammation-centered perspective while analyzing the networks constructed to understand VCI.

*Systems biology* or *network medicine* uses graph theory to analyze complex biological systems[8]. Protein-protein (PPI) interaction networks are used widely to analyze these complex biological systems for various diseases. Additionally, these PPIs can be modified to contain more information by enriching the nodes with their structures. Nevertheless, most studies construct static PPI networks such that each node represents one protein conformation. However, proteins are dynamic and can adopt many conformations, and each conformation they adopt can change the interactions they make and, therefore, the message they convey[9]. Depending on the conformation a protein adopts, it can interact differently with other proteins or perform different functions. These differential interactions can influence disease progression due to cellular functions, pathways, and signaling changes. Additionally, by finding out which interacting structures can be related to our disease of interest, we can investigate the possible effect of a mutation in that structure on the interaction it makes. As a result, the various interactions that PPI network nodes can make within the network can be revealed by enriching them with their alternative conformations. The interactions of proteins with their structures can be analyzed with molecular modeling tools like PRISM[10], allowing one to obtain deeper insights regarding the interactions of proteins.

VCI is a disease heavily influenced by inflammation and is gaining more and more attention. Nevertheless, there still is a considerable absence of knowledge of the molecular mechanisms related to it. Our work addresses this issue by analyzing the two most integral conditions related to it, CVD and CD, by constructing their related PPI networks. We merged these networks and analyzed the top three most connected clusters of the merged network as they contain insights regarding how CVD can affect CD and, therefore, VCI. We utilized the previously constructed networks for this ([11, 12]). We enriched these three clusters with their alternative conformations and used PRISM to predict the interactions between the conformations. We analyzed those interactions and mapped the mutations to those conformations for possibly annotated disease-related mutations. We found that chemokine-related and stress response-related pathways were enriched, possibly related to BBB dysregulation in VCI [11]. Our previous work also suggested the relevance of stress relted pathways and their effect in BBB. We suggested putative VCI-related interactions such as NFKBIA-RELA and proteasome complex and their effects. We found which mutated conformations’ interactions can be affecting VCI and that the following five interactions: LTF-SNCA, FGA-LTF, UBE2D1-VCP, ERBB4-INS, NFE2L2-VCP have a higher predicted binding affinity when one of the conformations are mutated. Lastly, we suggested that a cancer-related protein (BRCA1) could have implications in VCI as it interacts with VCP, which has a mutation in its conformation related to dementia. BRCA1 also interacts with mutated and wild-type XRCC4 and LIG4, implying the importance of the DNA damage response pathway, which might be shared between cancer and VCI. Altogether, our results suggest various pathways and interactions that can act as targets for therapeutic interventions or early diagnosis of VCI. These results can imply critical insights regarding VCI.

## RESULTS

As our previous analysis suggested that the inflammation-related proteins can have a critical role in the crosstalk of CVD and CD [11, 12], here we analyze their possible role in VCI. This study focuses on the role of inflammation in VCI by finding the inflammatory proteins in these global networks (GNs) that we previously constructed. We found 409 inflammatory proteins in our GNs, called core proteins (because we extended the PPI networks around them). Our network contains the interactors of the core proteins from the GNs and GWAS-related genes and, in total, resulting in 1728 genes. And the PPI network with these genes sums up to 1274 proteins and 87020 interactions (See methods section I-II). First set of results contains this PPI network’s top three most connected clusters and their enrichment with their structures (See methods section II-III). Molecular modeling with PRISM to analyze the structural PPI networks allowed us to analyze modeled interactions and whether mutated structures are interacting (See methods section IV). We demonstrated this workflow in **Figure 5**.

### Most connected clusters and Structural PPI Networks

Construction of a PPI network as inflammatory proteins being in the core, allowed us to analyze their potential role in VCI. As it would be computationally costly to enrich all the structures of such an extensive network, we advanced with the most connected three clusters of this network (**Table S1**). **Table S1** demonstrated that each cluster, from the most connected to the least connected, is rich in inflammatory-related proteins. The proteins present in these clusters are demonstrated in **Figure S1**. The cluster 1 (**Figure S1a**) is connected such that every node has a connection to every other node. Highly connected clusters are likely to be functionally significant, potentially representing essential biological modules within the network, making it crucial to perform functional enrichment on these three clusters(**Table 1**).

**Table 1.** Functional Enrichment of the three most connected clusters.

| Pathway/Cluster | Cluster 1 | Cluster 2 | Cluster 3 |
| --- | --- | --- | --- |
| GO:MF | chemokine activity<br>(GO:0008009) | lipoprotein particle binding<br>(GO:0071813) | identical protein binding<br>(GO:0042802) |
| GO:BP | chemokine-mediated signaling pathway<br>(GO:0070098) | inflammatory response<br>(GO:0006954) | response to stress<br>(GO:0006950) |
| GO:CC | extracellular space<br>(GO:0005615) | protein-lipid complex<br>(GO:0032994) | DNA repair complex<br>(GO:1990391) |
| KEGG | Viral protein interaction with cytokine and cytokine receptor<br>(KEGG:04061) | Alzheimer disease<br>(KEGG:05010) | Coronavirus disease - COVID-19<br>(KEGG:05171) |
| REAC | Chemokine receptors bind chemokine<br>(R-HSA-380108) | Interleukin-1 signaling<br>(R-HSA-9020702) | Response of EIF2AK4<br>(GCN2) to amino acid deficiency<br>(R-HSA-9633) |

The structural PPI network that was constructed is useful to model and analyze the interactions present in these clusters. However, unlike the traditional way of creating a structural PPI network by enriching one conformation per protein, we enrich each protein with its many alternative conformations. Each conformation represents one clustered group (as explained in the methods).

### Protein Modeling with PRISM

The interactions of the structural PPI network are modeled using PRISM (**Figure 1**). For the first cluster, there were four genes with missing UniProt IDs **(Table 3)**, and 10 and 13 for the second and third clusters. The remaining 21 proteins of the first cluster had 210 PPIs, and PRISM predicted 135 of these. Additionally, for the second and third clusters out of 155 and 131 PPIs, PRISM predicted 94 and 102 interactions, respectively. **Table 2** below demonstrates the top five least binding scores for each cluster. The binding score spans between −68.17 and −5.57 kcal/mol for cluster 1, −54.08 and −5.49 kcal/mol for cluster 2, and −36.05 and −5.30 kcal/mol for cluster 3. Furthermore, as knowing which specific conformations two proteins can interact with can provide valuable information, the PDB IDs of the interacting pairs are also provided.

**Figure 1.**
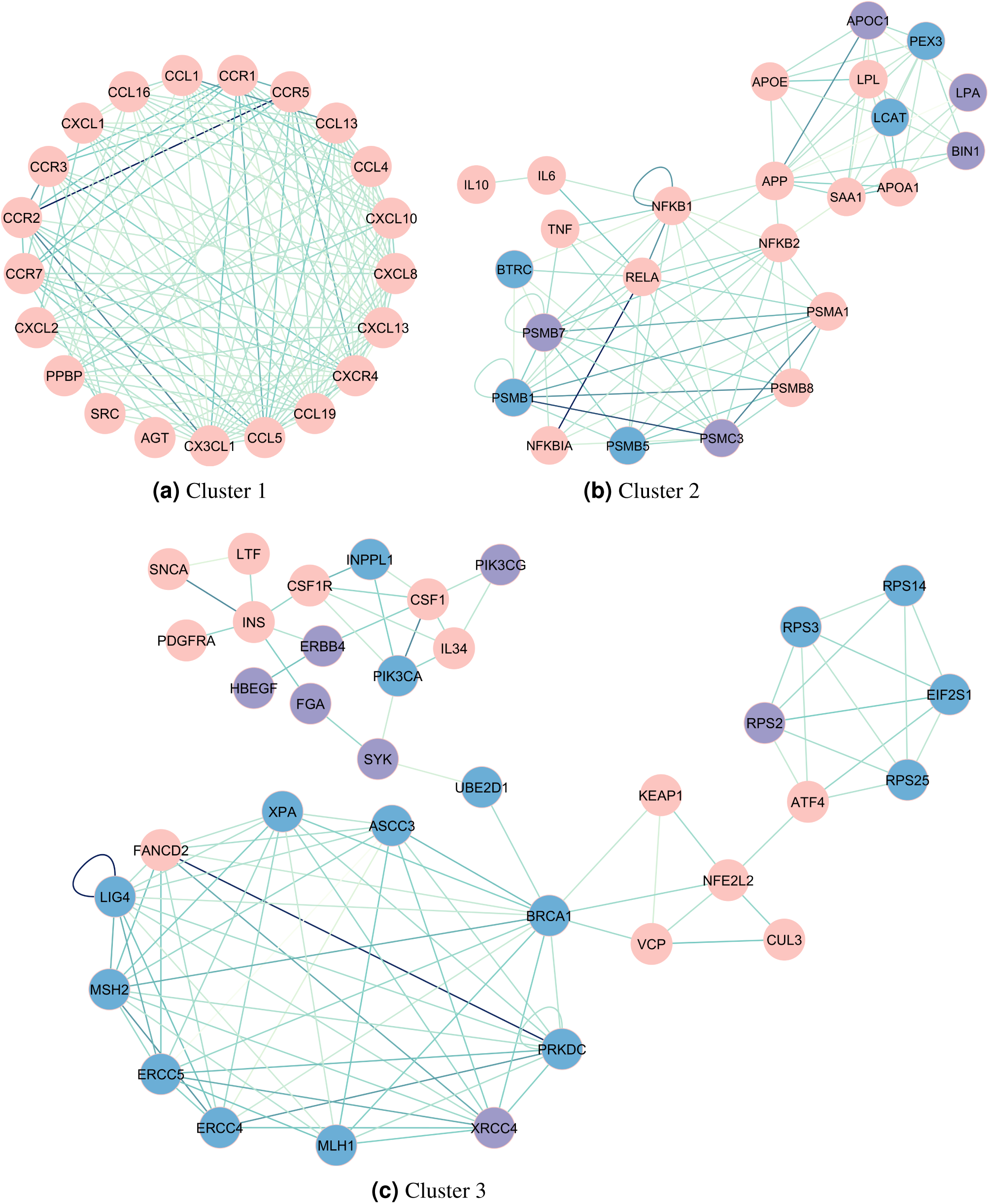
The PRISM modeled interactions ofr all three clusters. Darker edges represent lower PRISM binding score. Pink nodes represent core, purple nodes represent GWAS-related, and blue nodes represent interactor proteins. Self-edges occur when PDB-chain pair of a protein is interacting.

**Table 2.** Top five lowest predicted PRISM binding scores (B.S.) per cluster. The subscripts represent the chain of the interacting PDB.

| Cluster1 |  | Cluster2 |  | Cluster3 |  |
| --- | --- | --- | --- | --- | --- |
| Pair | B.S. | Pair | B.S. | Pair | B.S. |
| CCR2 - CCR5<br>5T1A<sub>A</sub> - 4MBS<sub>A</sub> | -68.17 | NFKBIA - RELA<br>1NFI<sub>C</sub> - 1IKND<sub>A</sub> | -54.08 | LIG4 - LIG4<br>3II6<sub>A</sub> - 3II6<sub>Y</sub> | -36.05 |
| CCR2 - CX3CL1<br>5T1A<sub>A</sub> - 7XBX<sub>R</sub> | -49.19 | PSMC3 - PSMB1<br>5M32<sub>G</sub> - 4R67<sub>A</sub> | -47.71 | PRKDC - FANCD2<br>7NFC<sub>K</sub> - 6VAA<sub>B</sub> | -35.15 |
| CCR2 - CCR3<br>7X9Y<sub>R</sub> - 7XA3<sub>R</sub> | -46.78 | PSMC3 - PSMA1<br>5M32<sub>G</sub> - 6KWY<sub>E</sub> | -39.72 | CSF1 - PIK3CA<br>4YKN<sub>A</sub> - 5LXF<sub>B</sub> | -26.79 |
| CCL1 - CCL13<br>2RA4<sub>A</sub> - 4OIK<sub>A</sub> | -45.81 | PSMB1 - PSMB8<br>5L5B<sub>L</sub> - 5L5A<sub>K</sub> | -38.04 | INS - SNCA<br>7RZF<sub>B</sub> - 1XQ8<sub>A</sub> | -26.38 |
| CCL5 - CCR2<br>7F1R<sub>R</sub> - 5T1A<sub>A</sub> | -41.27 | NFKB1 - RELA<br>2O61<sub>B</sub> - 1NFI<sub>C</sub> | -37.84 | MSH2 - ERCC4<br>8OLX<sub>A</sub> - 2MUT<sub>B</sub> | -25.24 |

**Table 3.**
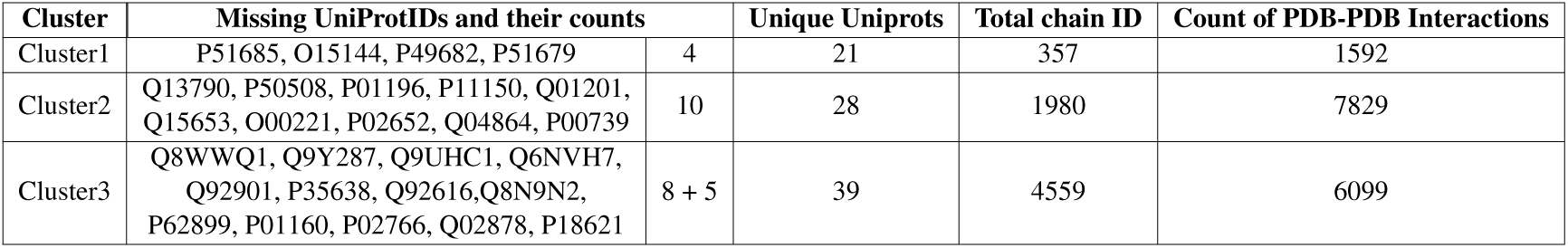
Table of UniProts. This table represents the proteins that do not have structures in PDB. For cluster 3, there are 5 additional uniprots at the end, in which the structure format is not in PDB.

| Cluster | Missing UniProtIDs and their counts |  | Unique Uniprotids | Total chain ID | Count of PDB-PDB Interactions |
| --- | --- | --- | --- | --- | --- |
| Cluster1 | P51685, O15144, P49682, P51679 | 4 | 21 | 357 | 1592 |
| Cluster2 | Q13790, P50508, P01196, P11150, Q01201, Q15653, O00221, P02652, Q04864, P00739 | 10 | 28 | 1980 | 7829 |
| Cluster3 | Q8WWQ1, Q9Y287, Q9UHC1, Q6NVH7, Q92901, P35638, Q92616, Q8N9N2, P62899, P01160, P02766, Q02878, P18621 | 8 + 5 | 39 | 4559 | 6099 |

### Mutated protein structures in the predicted interactions

We investigated the mutations for each interaction of the predicted protein interactions. None of the structures present in the first cluster had mutations. For the second cluster, SAA1, APP, and NFKB1 structures had many mutations. These mutated proteins made 11 interactions with each mutated and wild type (WT) structure. Moreover, lastly, INS, LIG4, LTF, PDGFRA, VCP, and XRCC4 had mutations in their structures. These made 24 interactions with each of their mutated and WT interactions (**Figure 2**)—the mutations these proteins have and their locations are given in **Table S2**

**Figure 2.**
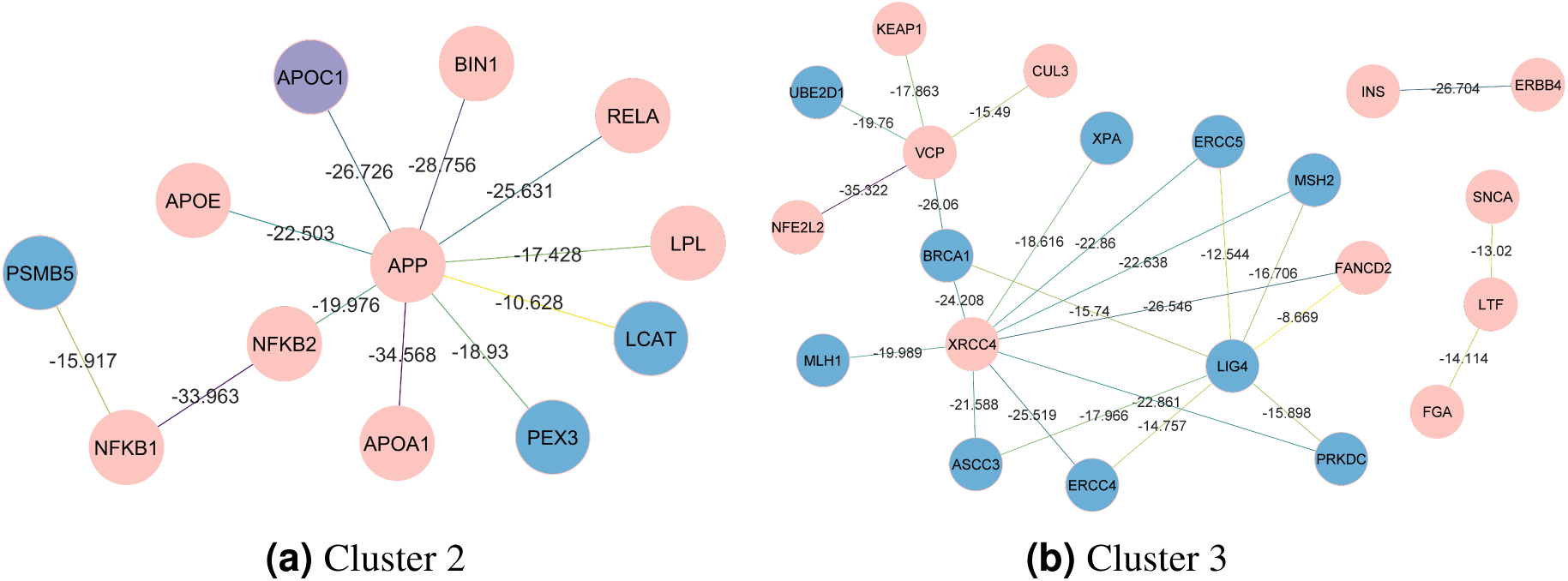
Each of these interactions represent that there is a mutated and wild type structure between the interacting proteins. The edges are colored such that darker colors represent lower binding score and thus higher binding affinity. The labels on the edges represent the predicted PRISM binding scores for the mutated and wild type structures interacting.

Each of these proteins participating in these interactions has both WT-WT and mutated-WT conformations interacting; each edge also has a predicted WT-WT binding score. The predicted binding score for the WT-WT conformations is referred to as *BS_nonmut_*, and the binding score between WT-mutated conformations is referred to as *BS_mut_*. We calculated the difference between these two scores (Δ*BS* = *BS_nonmut_ −BS_mut_*) to observe how the binding score changes when one of the protein conformations is mutated (**Supplementary Figure S2**). For cluster 2, two interactions (NFKB1-NFKB2 and NFKB1-PSMB5) have a positive Δ*BS*, indicating that the binding affinity increases when one of the conformations is mutated. The remaining nine interactions of cluster 2 from 2a have a negative Δ*BS*, indicating that the WT conformations have a higher binding affinity. On the other hand, five interactions of cluster 3 (LTF-SNCA, FGA-LTF, UBE2D1-VCP, ERBB4-INS, NFE2L2-VCP) have a positive Δ*BS*, and 19 have a negative Δ*BS*.

**Figure 2b** show interactions wherein breast cancer 1 (BRCA1) interacts with these three proteins’ non-mutant and mutated conformations (VCP, XRCC4, and LIG4). The **Figure 3** demonstrates the specific conformations predicted to interact in these PPIs.

**Figure 3.**
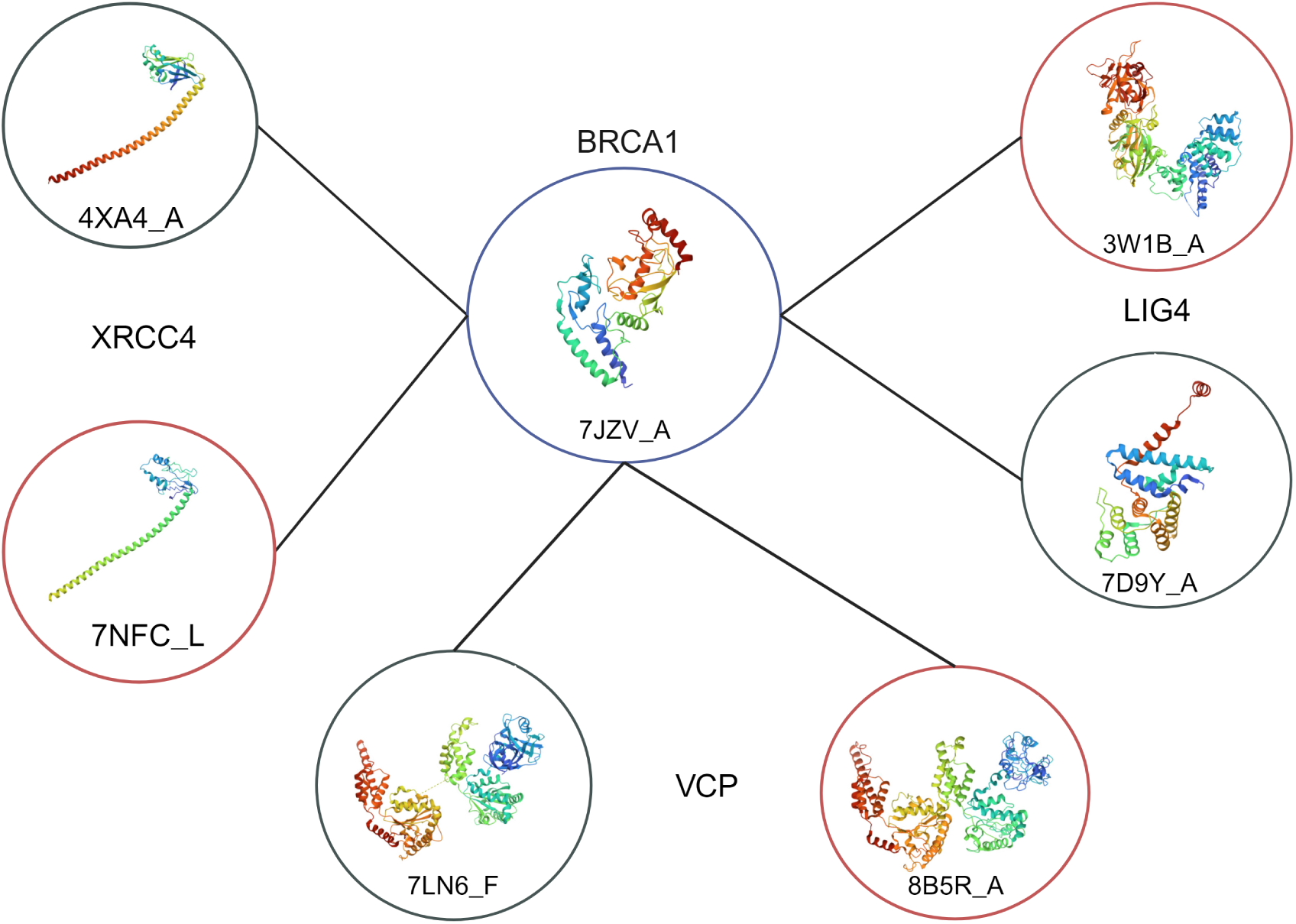
The mutated and wild type interactions BRCA1 makes according to PRISM predictions. Black circles represent mutated conformations and orange represents wild type conformations. The BRCA1 conformation is also nonmutated.

The interactions of BRCA1 with VCP, XRCC4, and LIG4 are closely investigated. According to PRISM modelling with lowest binding score, BRCA1 interacts with these interacting partners through the same region in both WT-WT and mutated-WT interactions. **Figure 4** show interactions of BRCA1 with VCP, XRCC4, and LIG4. BRCA1 interacts with BRCA1 interacts with VCP^WT^ with −30.45 binding score, whereas it interacts with VCP^MUT^ −23.25. BRCA1-XRCC4^WT^ interaction has a binding score of −27.72 and BRCA1-XRCC4^MUT^ interaction has a binding score of −17.36. Finally, the binding scores for the BRCA1-LIG4^WT^ and BRCA1-LIG4^MUT^ interactions are −18.14 and −15.74, respectively. Therefore, it is predicted that WT conformation of all VCP, XRCC4, and LIG4 with a higher binding score with respect to the mutated conformations, suggesting that WT-WT interactions are favoured.

**Figure 4.**
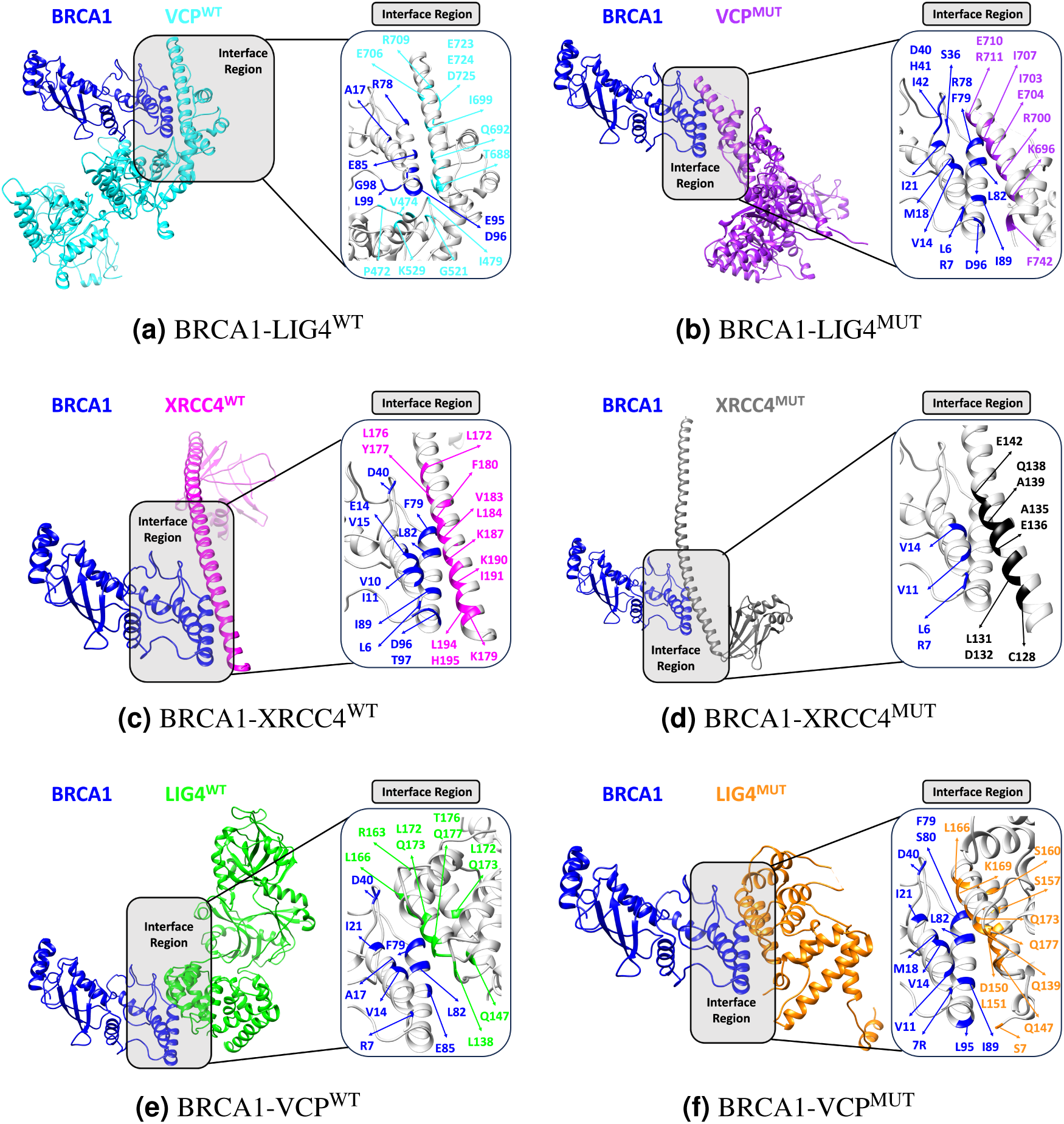
The interactions of BRCA1 with VCP, XRCC4, and LIG4 and the binding interfaces. Both WT-WT and mutated-WT interactions are shown

**Figure 5.**
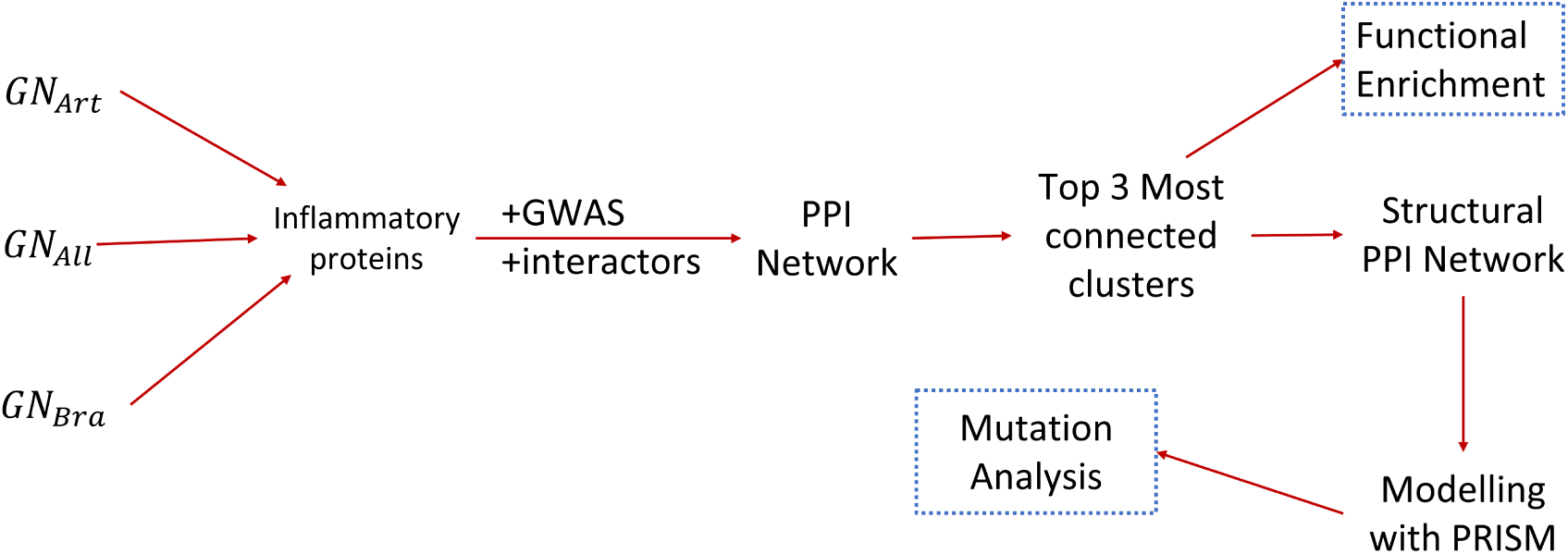
Workflow of the methods applied in the study.

BRCA1 is predicted to bind to VCP, XRCC4, and LIG4 through the same region in both WT-WT and mutated-WT interactions (**Supplementary Table S3**). Common interacting residues of BRCA1 for BRCA1-VCP^WT^ and BRCA1-VCP^WT^ interactions are L6, M18, R78, and D96. BRCA1 interacts with both XRCC4^WT^ and XRCC4^MUT^ with L6, V11, and V14. In both BRCA1-LIG4^WT^ and BRCA1-LIG4^MUT^ interactions BRCA1 interacts using the following residues: R7, V14, M18, I21, D40, R78, F79, and L82.

## DISCUSSION

Inflammation has a critical role in the progression of VCI. We discovered that pathways linked to the stress response and chemokines were enriched, which may be related to BBB dysregulation in VCI. We proposed potential VCI-related interactions and their consequences, including the NFKBIA-RELA and proteasome complex. The following five interactions: LTF-SNCA, FGA-LTF, UBE2D1-VCP, ERBB4-INS, and NFE2L2-VCP have a higher predicted binding affinity when one of the conformations is mutated. We discovered which mutated conformations’ interactions can affect VCI. Finally, we proposed that BRCA1 may play a role in VCI because it interacts with VCP, which possesses a conformational mutation linked to dementia. Additionally, BRCA1 interacts with both wild-type and mutated forms of XRCC4 and LIG4, suggesting the significance of the DNA damage response pathway, which may be common to both VCI and cancer. Collectively, our findings indicate multiple pathways and interactions that can act as targets for therapeutic interventions or early diagnosis of VCI.

### Inflammatory pathways related to VCI

The constructed PPI network and the most connected three clusters can provide essential insights regarding the role of the inflammatory proteins in VCI. The first cluster is a connected network, meaning that every protein is making all the possible connections they can make (**Table S1**). Additionally, from the first to the third cluster, while the count of core proteins decreases, the count of interactor and GWAS-related proteins increases. From this, we can deduce that the core of the constructed inflammatory-centered network does contain inflammatory-related proteins on its core. The presence of the interactor and GWAS-related proteins on the other most connected clusters supports the importance of creating an extensive PPI network.

Finding the most connected clusters in a network is vital because it serves the purpose of finding functionally relevant biological modules [13]. According to the functional enrichment of these clusters (**Table 1**), the genes in the first cluster are enriched in chemokine-related activity. This activity suggests the specific importance of chemokines in VCI. Moreover, chemokines assist the migration of monocytes and lymphocytes(inflammatory proteins) across the endothelial lining of blood arteries[14]. This migration process is essential in pathological conditions like Alzheimer’s disease (AD) [15], atherosclerosis [16]. Moreover, neurons [17] and endothelial cells [18] have chemokine receptors on their surface. The malfunction of these receptors can thus be critical regarding the dysregulation of the blood-brain barrier (BBB) as endothelial cells are also present in the BBB[19]. Therefore, as the BBB is critical for cognitive health, its dysregulation could significantly contribute to the pathophysiology of VCI.

The second most connected clusters’ functional enrichment infers that the disruptions in lipid metabolism and inflammatory processes can be critical. The dysregulation of the lipid metabolism was linked to increased inflammatory response and excess in circulating fatty acids, thus affecting the cognitive function[20, 21, 22]. Further, the lipid transport and metabolism are essential for preserving the integrity of neuronal membranes[23] and can, therefore, contribute to the pathophysiology of VCI. Amyloid-beta metabolism and tau pathology are known to be related to AD [24]. The enrichment of AD in KEGG pathways can suggest that these mechanisms are overlapping in AD and VCI. IL-1, on the other hand, is a pro-inflammatory cytokine related to neurodegeneration, neuronal survival, and synaptic plasticity [25, 26].

The third cluster is enriched with more diverse functions. Stress responses can cause BBB malfunction and vascular alterations[27]. On the other hand, response of EIF2AK4 (GCN2) to amino acid deficiency is related protein synthesis, which is essential for the cell’s survival under metabolic stress[28]. Previous research has linked dysfunctions in the DNA repair pathways to aging and neurodegenerative diseases[29]. Enrichment of the DNA repair complex underscores the importance of genomic integrity for neuronal survival and function and, therefore, cognitive health[30, 31]. Studies reported that some viral infections (like cytomegalovirus (CMV), or COVID-19) have associations with vascular health and systemic inflammation [32, 33]. Interestingly, the enrichment of COVID-19 could emphasize the effects of these implications and their possible burden on cognitive function. Moreover, there is an increasing amount of ongoing research regarding the effect of COVID-19 on the central nervous system and its possible effect on cognitive decline [34, 35, 36].

All of the discussed pathways contribute to understanding VCI. These pathways provide a central view regarding the roles of the inflammatory-related pathways in this crosstalk. More research on these pathways concerning VCI is required to better understand the mechanisms behind these disorders.

### Putative VCI related interactions

Performing molecular modeling with PRISM to every conformation pair for each interaction in each cluster resulted in many predictions. PRISM predicted most of the PPIs. We investigated the top five interactions for each cluster that has the highest predicted binding affinity (**Table2**). While it is crucial to know which of these proteins and interactions were previously related to VCI, we also surveyed the literature on whether the specific conformations were mentioned to be related to VCI pathophysiology. In the first cluster, CCR2 made four of the top five most substantial interactions regarding the predicted binding score. CCR2 is the CC chemokine receptor 2. The interaction between this receptor and its ligand (CCL2/ monocyte chemoattractant protein-1, MCP-1) is essential in the monocyte migration and infiltration into the brain[37, 38]. Studies have shown that dysregulation in CCR2 could worsen cognitive deficits in mice models[39, 40]. Studies demonstrated that the heterodimer formation of CCR2 and CCR5 could lead to differential coupling with other proteins, causing changes in signal regulation [41, 42]. Additionally, Mellado et al. suggested that this heterodimerization may be favored by low chemokine concentrations[43]. CX3CL1 (Fractalkine) is a chemokine expressed by brain neurons. It mainly regulates microglial activation and neuroinflammation by interacting with its receptor (CX3CR1). The relationship between this ligand and its receptor was linked to cognitive health by Rogers et al. [44]. In mice models, a deficit of CX3CL1 showed a significant decline in cognitive function[45]. To our knowledge, the interaction between CCR2-CCR3 and CCR2-CX3CL1 is not known in the current literature. Nevertheless, these interactions seem plausible because of the ability of heterodimer formation between CCR2-CCR5 and due to the crosstalk of the CCR2-CX3CR1 pathways. These specific interactions of the first cluster given in **Table 2** were not previously proposed to be VCI-related. We further propose that the given conformations for each interaction can be important therapeutic targets.

Regarding the second cluster, to our knowledge, none of these interactions and specifically interacting conformations are directly related to VCI. The pathways connected to inflammation and the immune response include NFKBIA-RELA. In the NF-KB signaling pathway, NFKBIA is an inhibitory molecule that combines with RELA (p65). RELA is a critical component of the NF-KB signaling pathway, linked to several inflammatory diseases like multiple sclerosis and atherosclerosis [46]. Furthormore, research indicated that NFKBIA could be a biomarker in diagnosing Alzheimer’s Disease[47]. The interactions in the second cluster are also related to the proteasome complex due to the PSMA, PSMB, and PSMC proteins. Dysregulation in the proteasome complex can cause neurodegenerative diseases. The proteasome complex is essential in preventing protein degradation, and when this mechanism does not work as it should, the resulting protein accumulation can cause neurodegenerative diseases. The proteasome complex is also involved in regulating the NF-KB signaling pathway. Our results indicate that the dysfunction in the crosstalk of these pathways can exacerbate the neuroinflammation, contributing to the pathophysiology of VCI. In the context of the third cluster, one of these interactions or conformations was also related to VCI. These interactions, nevertheless, highlight the importance of DNA repair mechanisms, as discussed in the previous section.

### Conformational preferences of the mutated inflammatory proteins

#### Increased affinity favoring mutated conformations in five interactions

Our results suggested that five interactions have a positive Δ*BS*; this indicates that the predicted PRISM binding affinity increases when the conformation is mutated. These five interactions are **LTF**-SNCA, FGA-**LTF**, UBE2D1-**VCP**, ERBB4-**INS**, NFE2L2-**VCP** (The bold ones are the proteins that have a mutated conformation interacting with the other nonmutated protein.).

Two of these interactions contain VCP (valosin-containing protein) which is a calcium-associated ATPase protein functions in ER and mithochondria. This protein plays a vital role in controlling protein aggregates, calcium homeostasis and the ubiquitin-proteasome pathway, both essential for vascular health and neurodegeneration[48]. The mutations of this protein were related to Frontotemporal Dementia and/or Amyotrophic Lateral Sclerosis 6 (FTDALS6) and Inclusion Body Myopathy with Paget’s Disease of Bone and Frontotemporal Dementia (IBMPFD1) [49, 50]. VCP’s interaction with UBE2D1, an enzyme that conjugates ubiquitin, emphasizes the function of the ubiquitin-proteasome system [51]. Additionally, VCP’s interaction with nuclear factor erythroid 2–related factor 2 (NFE2L2), a crucial regulator of the antioxidant response, highlights the role that oxidative stress defenses play in preserving the health of neurons [52]. These pathways are vital in mitigating protein misfolding and oxidative damage; vascular complications aggravate these consequences and may result in cognitive impairments. Our previous results also supported the importance of oxidative stress in VCI pathophysiology.

Fibrinogen(FGA) is a blood coagulation factor implicated in vascular disease and here interacts lactotransferrin (LTF /LF, lactoferrin). Previous research suggested that FGA contributes to cerebrovascular dysfunction and that this dysfunction is linked to cognitive decline [53]. Lactotransferrin is a multifunctional protein of the transferrin family. It controls the immune system and has a role in iron transport[54]. Additionally, a mutation in the structure of LTF is related to coronary artery stenosis [55]). This interaction emphasizes the interaction between coagulation and inflammatory response and further its possible effect on VCI pathology. The receptor tyrosine kinase Erb-B2 Receptor Tyrosine Kinase 4 (ERBB4) is involved in the development of neurons and the plasticity of synapses[56]. Insulin (INS) is a crucial regulator of glucose metabolism. This interaction may provide a relationship such that metabolic elements, controlled by insulin signaling, are essential in VCI.

Alpha-synuclein (SNCA) and LTF have an intriguing interaction because SNCA is linked to neurodegenerative diseases like Parkinsons disease[57]. These two proteins are intercting with medium confidence according to STRINGdb score. Furthermore, LTF provides a vascular component to this interaction since a mutation in the structure of LTF was related to coronary artery stenosis. Therefore, the relationship between LTF and SNCA may be a crucial point where vascular inflammation due to LTF mutations, may affect SNCA aggregation and contribute to the cognitive decline associated with VCI. Here, we propose that this interaction could be significant in the vascular-cognitive health axis.

By identifying five PPIs (LTF-SNCA, FGA-LTF, UBE2D1-VCP, ERBB4-INS, and NFE2L2-VCP), we provided significant new insight into the intricate pathophysiology of VCI. These interactions sheds light on the complex molecular network underlying the neurodegenerative and vascular components of VCI. Given that these proteins bind with more affinity when one of them is mutated, these changed interactions may exacerbate the pathogenic mechanisms causing cognitive decline. For example, vascular inflammation may be exacerbated by the mutation in LTF linked to coronary artery stenosis. Also, additional disruption of protein homeostasis may result from VCP mutations associated with neurodegenerative disorders, which could worsen the damage to neurons. Moreover, none of these mutations were previously related to VCI directly to our knowledge, and neither were the interactions they make. These results can be critical in understanding complex vascular factors that contribute to cognitive impairment and can provide specific therapeutic targets.

#### BRCA1 and its possible cancer relation to VCI

BRCA1 is a crucial protein in DNA repair and maintaining genomic stability[58]. This protein interacts with essential elements of the cellular stress pathways and DNA damage response[59, 60]. These interactions that BRCA1 makes may suggest an intricate network that may impact the pathophysiology of VCI and cancer progression.

LIG4 (DNA Ligase IV) is a protein that repairs double-strand DNA breaks. LIG4 forms a complex with a DNA repair protein X-ray repair cross-complementing protein 4 (XRCC4) [61]. Interestingly, the mutations associated with the conformations of these proteins were cancer-related and uncertain. Meanwhile, VCP is involved in protein degradation and cellular stress responses, as previously mentioned. Mutated forms of these three proteins interacting with BRCA1 imply disruptions in DNA damage response elements may be related to cancer pathogenesis by compromising DNA repair pathways, potentially leading to genomic instability, a hallmark of cancer development. Moreover, studies indicate that cellular stress (or OS) can cause genomic instability in the brain which in turn could play a role in neurodegenerative processes[62, 63]. This instability can happen by the chronic activation of the DNA damage response in neurons, resulting in many cognitive decline-related processes, such as cellular senescence, inflammation, and apoptosis[64, 65, 66]. In order to preserve the integrity of the vascular endothelium and prevent cerebrovascular disorders, genomic stability and effective DNA repair are crucial. Furthermore, there are mutations on the BRCA1 protein that were previously related to CVD[67], making BRCA1 a protein related to both CVD and CD but lakcking substantial suggesion in its role to VCI.

We further investigated the domains and sites in the interfaces of these structures. Regarding the interaction between LIG4 and BRCA1, the interacting residues of LIG4 fall into RING-type domain. In the interaction between XRCC4 and BRCA1, the interacting residues of XRCC4 for the mutated version normally interacts with IFFO, and for the WT version it is a RING-type domain. Mutation in this protein can be affecting its interaction with other proteins. And lastly for VCP, both of the conformations’ interacting sites are disordered regions. Disordered regions can be pointing out to dynamic nature of the interface, leading to many interactions and further continuation of interaction when protein is mutated.

Mutations in the DNA damage response system may influence vascular health and cancer susceptibility, which in turn raises the chance of cognitive impairment. Except for the mutations on VCP, none of the mentioned mutations were previously related to VCI. Our results suggest that these mutations in DNA damage response mechanisms can play a critical role in VCI and can also be related to cancer pathways. Additionally, to our knowledge, none of the structures were mentioned to be VCI-related in the literature. Here, we suggest that the interactions of these proteins and the conformations in which they interact can be crucial regarding the mechanisms behind VCI. Our results also suggest that more research is needed to determine the shared mechanisms of cancer and neurodegenerative diseases.

## CONCLUSIONS AND FUTURE DIRECTIONS

We found that chemokine-related, and stress response-related pathways were enriched, possibly related to BBB dysregulation in VCI. Due to malfunctions in lipid metabolism and DNA repair pathways, inflammatory response may increase in VCI. Viral infections that affect cognitive decline can also be a factor in VCI. We suggested putative VCI-related interactions such as NFKBIA-RELA and proteasome complex related and their effects. We found which mutated conformations’ interactions can be affecting VCI and that the following five interactions: LTF-SNCA, FGA-LTF, UBE2D1-VCP, ERBB4-INS, NFE2L2-VCP have a higher binding affinity when one of the conformations are mutated. This can imply critical insights regarding VCI. Lastly, we suggested that a cancer-related protein (BRCA1) could have implications in VCI as it interacts with VCP, which has a mutation in its conformation related to dementia. BRCA1 also interacts with mutated and wild-type XRCC4, and LIG4 implying the importance of the DNA damage response pathway, which might be shared between cancer and VCI. Altogether, our results suggest various pathways and interactions that can act as targets for therapeutic interventions or early diagnosis of VCI.

## METHODS

### Section I: Global Networks representing the CVD-CD crosstalk

We previously studied the proteins and pathways related to VCI by analyzing the crosstalk between CVD and CD. For this, previously, we constructed Global Networks (GNs) representing VCI using three different tissue settings [68][12]. We studied the crosstalk between CVD and CD by constructing PPI networks and selected the CVD and CDs most relevant to VCI. In our previous study, we developed a disease-phenotype taxonomy. We used GUILDify web server 2.0 [69] to find the diseases with the most seed proteins (highest relevance in literature). We selected Atherosclerosis, Atrial Fibrillation, Hypertension, Myocardial Ischemia, and Stroke for CVD, and Dementia and Impaired Cognition for CD. Furthermore, as the OS network is a crucial mechanism for the interaction between CD and CVD, we constructed it. We obtained the GN by combining these eight networks. In our previous work, we constructed the GN in the “all”, artery and brain tissue setting. In this study, we utilized all three of these GNs.

### Section II: Selection of inflammasome and inflammation-associated proteins and PPI Network Construction

We selected the proteins related to the inflammasome by searching “inflammasome” in UniProt and later filtering the results by “reviewed” and by “Homo Sapiens.” This resulted in 124 proteins. To encompass a broad range of proteins related to inflammasome and inflammatory system and also to better understand the role of this system, we also searched for “inflammatory” in UniProt. We later filtered the results by “reviewed” and by “Homo Sapiens.” This part resulted in 1217 proteins. We obtained a total of 1341 inflammatory-related proteins. There were 1263 unique inflammatory-related proteins. Four hundred nine of these inflammatory-related proteins were present in our GNs. These 409 core proteins had 709 interactors in the GNs.

We also included GWAS-related genes for the diseases we selected in order to extend the information that can be obtained from our PPI networks. We extracted GWAS data using the API of EBI. We extracted the GWAS genes of the following traits. Atherosclerosis: EFO 0003914, Stroke: EFO 0000712, Atrial fibrillation: EFO 0000275, Hypertension: EFO 0000537, Myocardial Ischemia: EFO 1001375, Dementia: HP 0000726, Cognitive Impairment: HP 0100543, Vascular dementia: EFO 0004718 [70]. After including these GWAS-related genes, we had 1728 genes in total.

We created a PPI network with these 1728 genes using STRINGdb[71], which resulted in a PPI network of 1589 genes and 30840 interactions. The reduction in the amount of genes is because of the long non coding rnas and open reading frame related genes. We downloaded the medium confidence (STRINGdb score of at least0.4) interactions known with experiments and databases. We analyzed this network using Cytoscape[72], which had 1589 nodes with 8702 interactions, and the most connected network here had 1274 nodes.

We used the MCODE[13] application of Cytoscape to find the most connected clusters of this PPI network. We used the default MCODE settings, which did not include loops and fluffs, with a degree cutoff of 2, node score cutoff of 0.2, K-core 2, and a maximum depth of 100. The MCODE scores of the three clusters are respectively as follows: 25, 15.78, 9.73. These three most connected clusters were used for the pathway enrichment and construction of structural PPI network.

### Section III: Structural PPI Network

To create the structural PPI of each cluster, we need all the structures in PDB related to that protein. However, as PDBs can be redundant, clustering similar PDB structures together is necessary. In this case, a protein can have alternative conformations such that each conformation is representative of that cluster. In other words, each cluster for a protein represents an alternative conformation of that protein, and each cluster is assigned a representative PDB. We did clustering through the use of sequence identity and structural similarity. We did not include proteins with fewer than 30 residues. When two PDB structures share at least 95% of their amino acid sequences and have an RMSD value less than 2 Angstroms, we consider them in the same cluster. Finally, we constructed the structural PPI network (PDB-PDB network) using a set of scripts and fed them into PRISM protein modeling. For this, we inspected the UniprotIDs and the PDBs of each protein. **Table3** below demonstrates the missing PDBs of the corresponding proteins UniprotID. For the PRISM part of this analysis, we will use each PDB-chain pair.

### Section IV: PRISM and Mutation Analysis

PRISM is a method that uses protein structural analysis and template-based modeling of interactions between proteins to predict these interactions [10]. PRISM predicts the interactions between each PDB-chain pair from the constructed structural PPI.

We also analyzed the possible mutations in these predicted interactions. We collected the mutation information from ClinVar [73] and UniProt [74]. For this, we mapped the mutations to the expected PRISM modelings to investigate the impact of the mutations on the structural PPI. This study solely took single nucleotide variations into account.

## ACKNOWLEDGMENTS.

This project has been funded in part by TUBITAK Research Grant No: 220N252 and European Research Area Net (ERANET) ERA-CVD JTC2020-015.

**SUPPORTING INFORMATION.**

## SUPPLEMENTARY INFORMATION

### FIGURES

**Figure S1.**
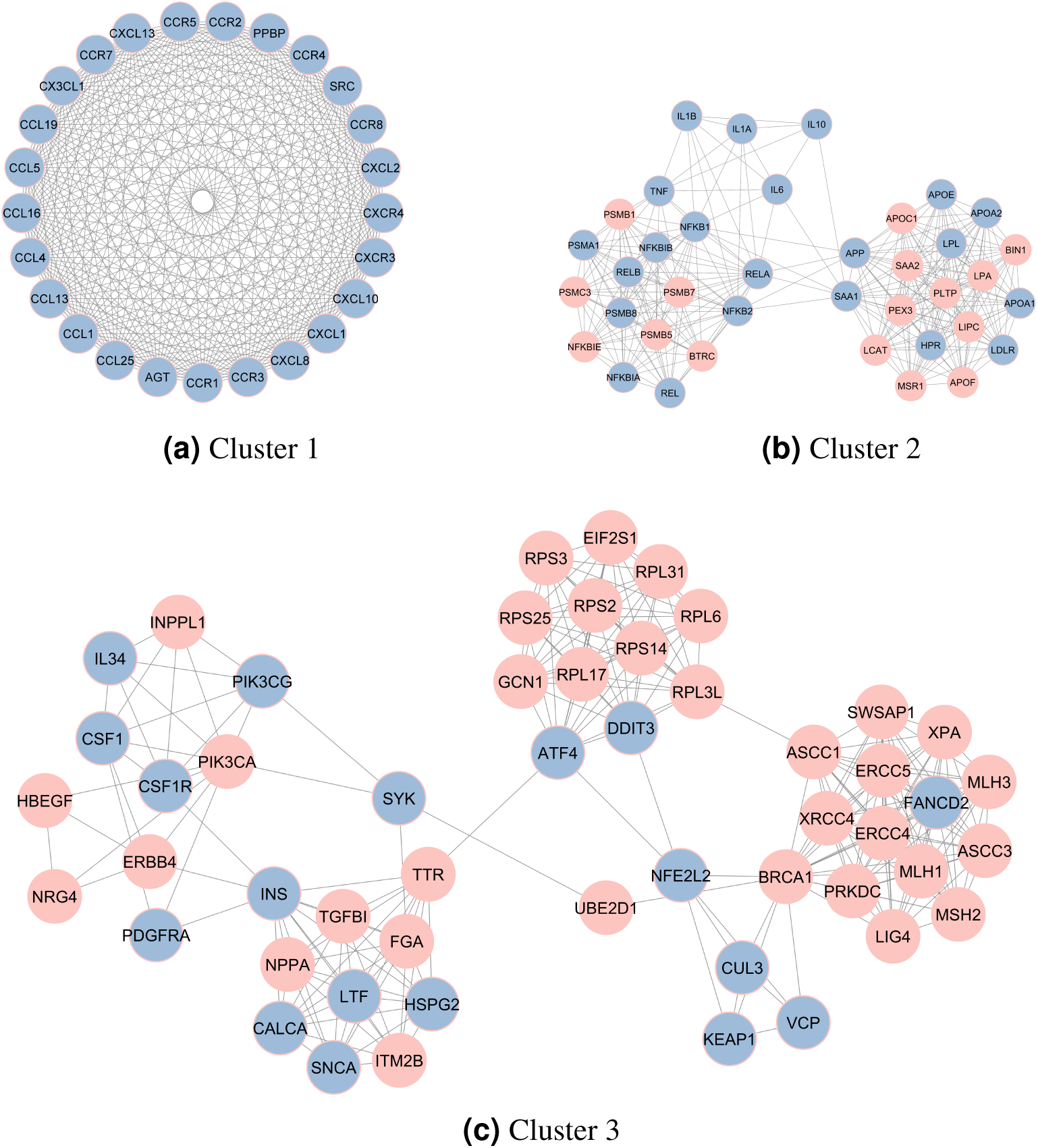
Top 3 clusters of the constructed networks. The pink nodes represent the core proteins (inflammatory-related proteins.

**Figure S2.**
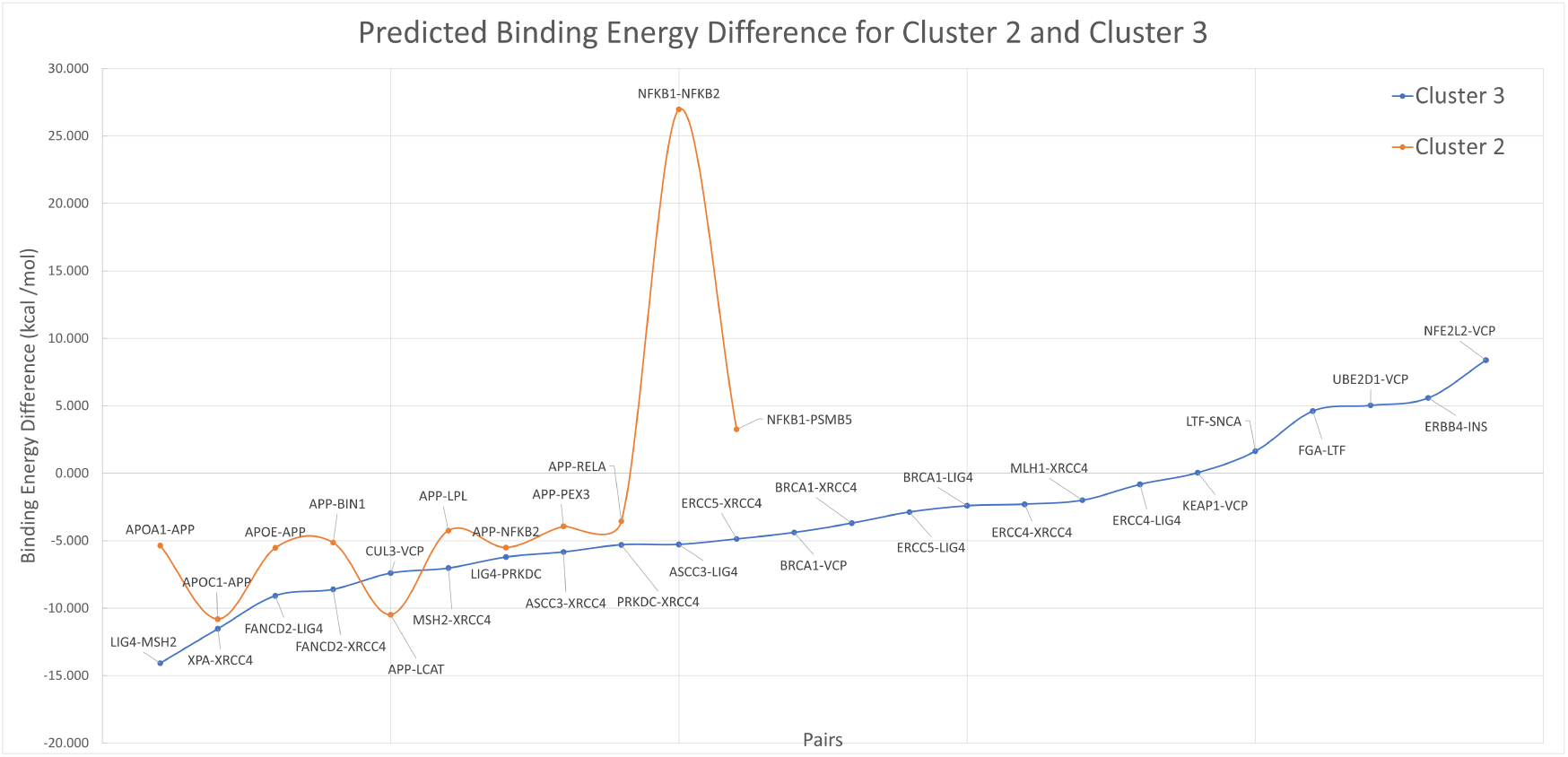
Difference between nonmutmut binding structures

### TABLES

**Table S1.**
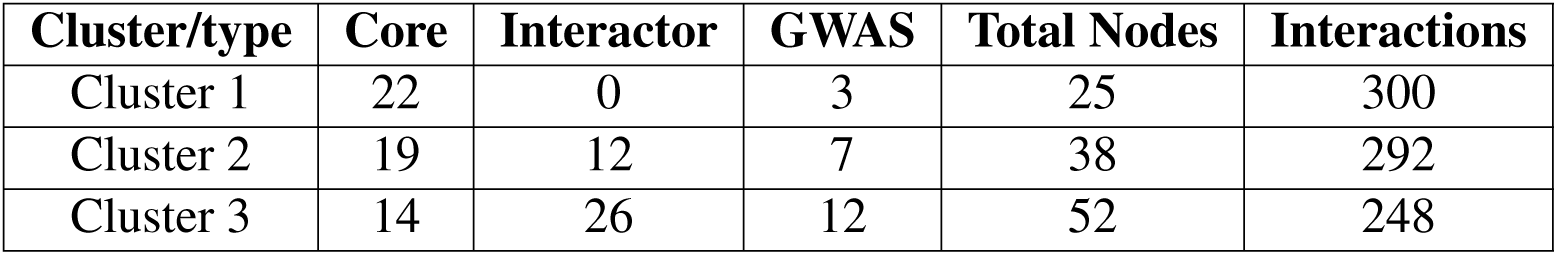
Top three most connected clusters and the count of core, interactor, GWAS-related proteins in those clusters.

**Table S2.**
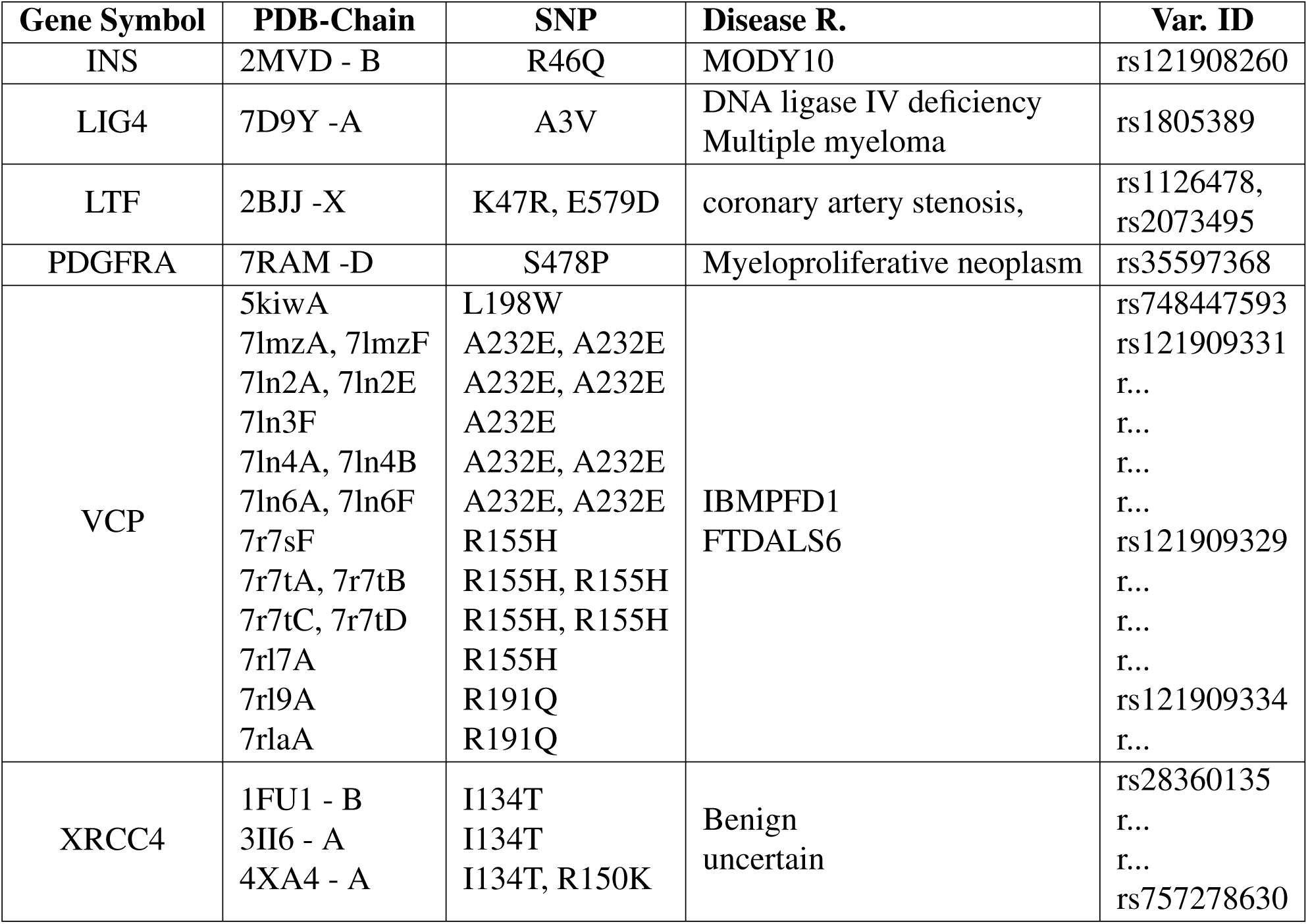
Mutations in the structural network. If the variant IDs are the as the previous variant ID it is shown as “r…”.

**Table S3.**
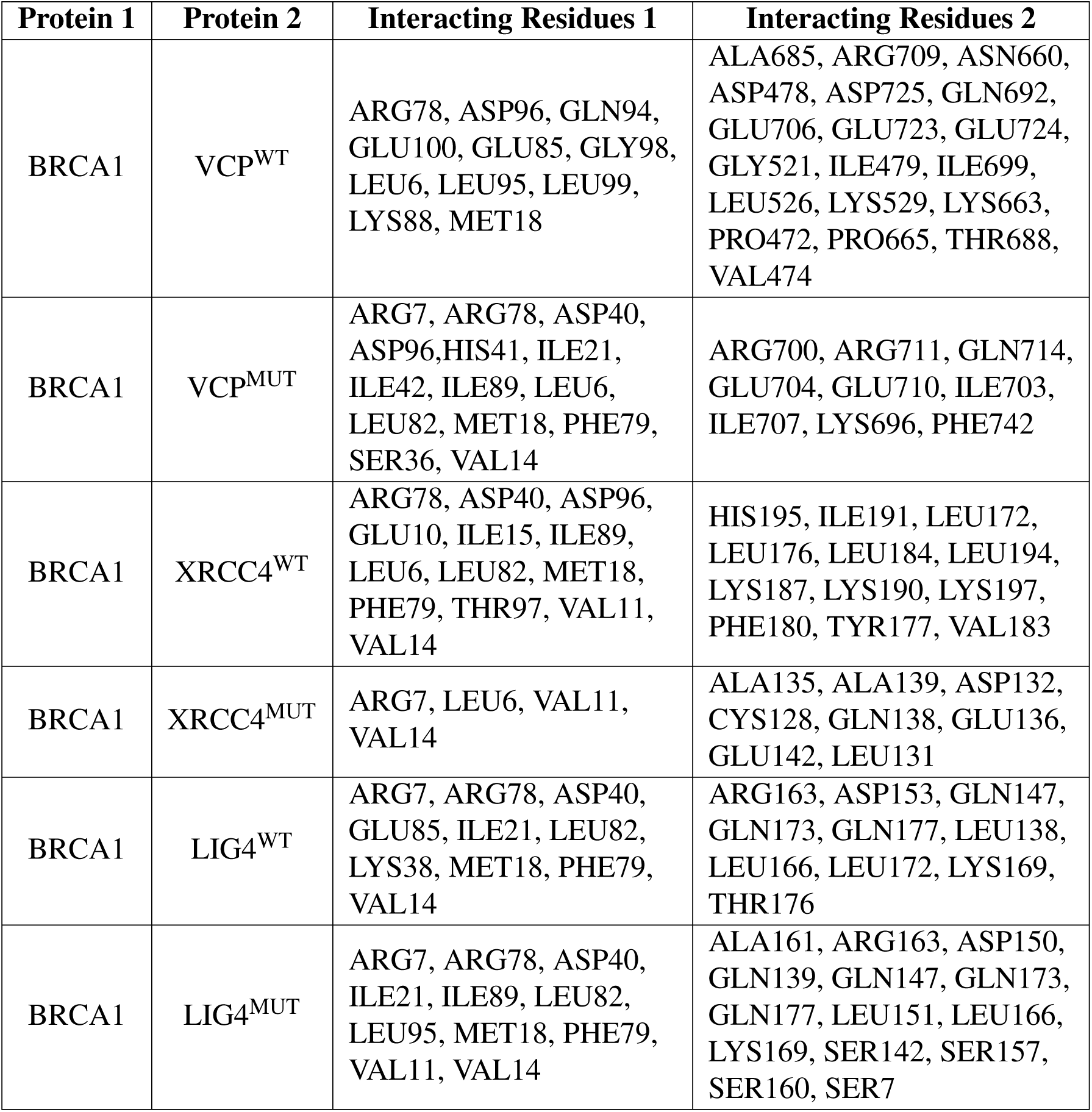
Interface residues of BRCA1-VCP, BRCA1-XRCC4, and BRCA1-LIG4 interactions.

## AUTHOR CONTRIBUTIONS

M.E.Z., S.S., A.G., and O.K. designed and conceptualized the project. M.E.Z. and S.S. analyzed data, prepared tables and figures. M.E.Z., S.S., A.G., O.K. wrote and edited the manuscript. All of the authors reviewed and approved the final manuscript.

## DATA AVAILABILITY

The datasets generated during and/or analyzed during the current study are available from the corresponding author on reasonable request.

## COMPETING INTERESTS

The author(s) declare no competing interests.

## Notes

### Competing Interest Statement

The authors have declared no competing interest.

