## Supplementary Information for "An Inflammation Centered Perspective to the Mechanisms and Interactions Related to Vascular Cognitive Impairment"

#### FIGURES

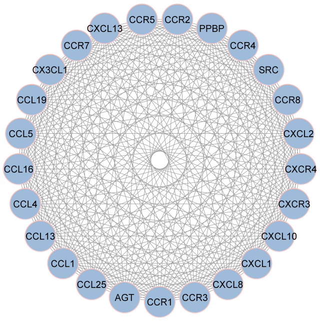


(a) Cluster 1

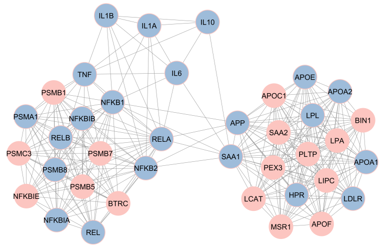


(b) Cluster 2

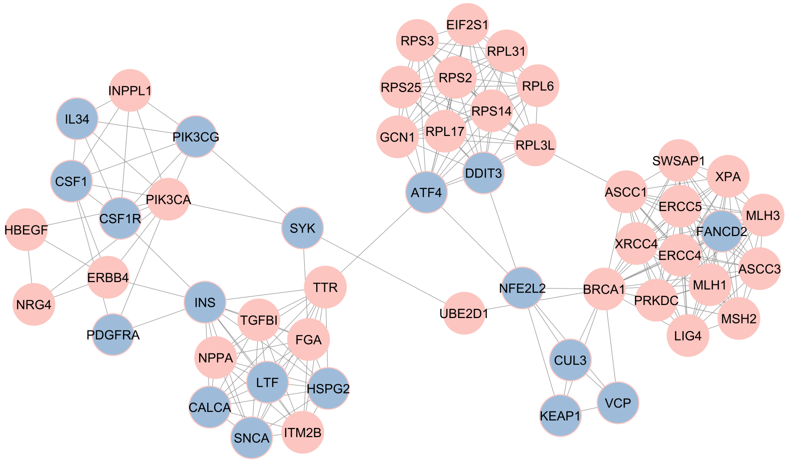


(c) Cluster 3

Figure S1: Top 3 clusters of the constructed networks. The pink nodes represent the core proteins ( inflammatory-related proteins.

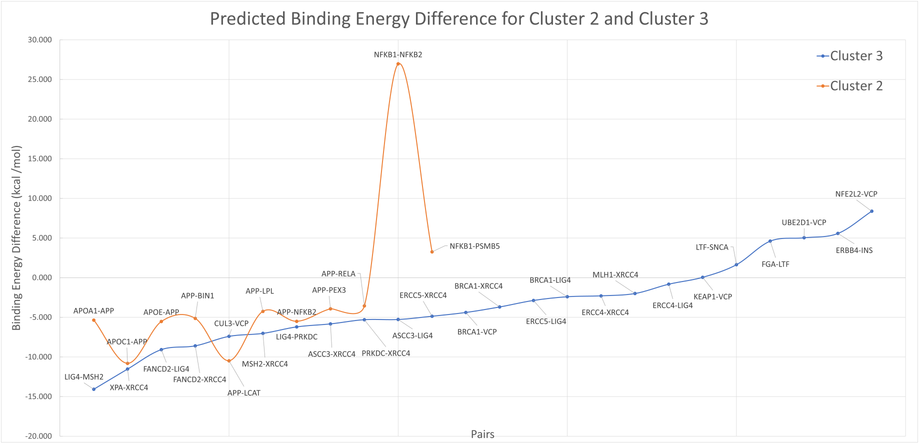


Figure S2: Difference between nonmut- mut binding structures

### TABLES

Table S1: Top three most connected clusters and the count of core, interactor, GWAS-related proteins in those clusters.

| Cluster/type | Core | Interactor | GWAS | Total Nodes | Interactions |
|---|---|---|---|---|---|
| Cluster 1 | 22 | 0 | 3 | 25 | 300 |
| Cluster 2 | 19 | 12 | 7 | 38 | 292 |
| Cluster 3 | 14 | 26 | 12 | 52 | 248 |

Table S2: Mutations in the structural network. If the variant IDs are the as the previous variant ID it is shown as ”r...”

| Gene Symbol | PDB-Chain | SNP | Disease R. | Var. ID |
|---|---|---|---|---|
| INS | 2MVD - B | R46Q | MODY10 | rs121908260 |
| LIG4 | 7D9Y -A | A3V | DNA ligase IV deficiency<br>Multiple myeloma | rs1805389 |
| LTF | 2BJJ -X | K47R, E579D | coronary artery stenosis, | rs1126478,<br>rs2073495 |
| PDGFRA | 7RAM -D | S478P | Myeloproliferative neoplasm | rs35597368 |
| VCP | 5kiwA<br>7lmzA, 7lmzF<br>7ln2A, 7ln2E<br>7ln3F<br>7ln4A, 7ln4B<br>7ln6A, 7ln6F<br>7r7sF<br>7r7tA, 7r7tB<br>7r7tC, 7r7tD<br>7rl7A<br>7rl9A<br>7rlaA | L198W<br>A232E, A232E<br>A232E, A232E<br>A232E<br>A232E, A232E<br>A232E, A232E<br>R155H<br>R155H, R155H<br>R155H, R155H<br>R155H<br>R191Q<br>R191Q | IBMPFD1<br>FTDALS6 | rs748447593<br>rs121909331<br>r...<br>r...<br>r...<br>r...<br>rs121909329<br>r...<br>r...<br>r...<br>rs121909334<br>r... |
| XRCC4 | 1FU1 - B<br>3II6 - A<br>4XA4 - A | I134T<br>I134T<br>I134T, R150K | Benign<br>uncertain | rs28360135<br>r...<br>r...<br>rs757278630 |

Table S3: Interface residues of BRCA1-VCP, BRCA1-XRCC4, and BRCA1-LIG4 interactions.

| Protein 1 | Protein 2 | Interacting Residues 1 | Interacting Residues 2 |
|---|---|---|---|
| BRCA1 | $VCP^{WT}$ | ARG78, ASP96, GLN94, GLU100, GLU85, GLY98, LEU6, LEU95, LEU99, LYS88, MET18 | ALA685, ARG709, ASN660, ASP478, ASP725, GLN692, GLU706, GLU723, GLU724, GLY521, ILE479, ILE699, LEU526, LYS529, LYS663, PRO472, PRO665, THR688, VAL474 |
| BRCA1 | $VCP^{MUT}$ | ARG7, ARG78, ASP40, ASP96,HIS41, ILE21, ILE42, ILE89, LEU6, LEU82, MET18, PHE79, SER36, VAL14 | ARG700, ARG711, GLN714, GLU704, GLU710, ILE703, ILE707, LYS696, PHE742 |
| BRCA1 | $XRCC4^{WT}$ | ARG78, ASP40, ASP96, GLU10, ILE15, ILE89, LEU6, LEU82, MET18, PHE79, THR97, VAL11, VAL14 | HIS195, ILE191, LEU172, LEU176, LEU184, LEU194, LYS187, LYS190, LYS197, PHE180, TYR177, VAL183 |
| BRCA1 | $XRCC4^{MUT}$ | ARG7, LEU6, VAL11, VAL14 | ALA135, ALA139, ASP132, CYS128, GLN138, GLU136, GLU142, LEU131 |
| BRCA1 | $LIG4^{WT}$ | ARG7, ARG78, ASP40, GLU85, ILE21, LEU82, LYS38, MET18, PHE79, VAL14 | ARG163, ASP153, GLN147, GLN173, GLN177, LEU138, LEU166, LEU172, LYS169, THR176 |
| BRCA1 | $LIG4^{MUT}$ | ARG7, ARG78, ASP40, ILE21, ILE89, LEU82, LEU95, MET18, PHE79, VAL11, VAL14 | ALA161, ARG163, ASP150, GLN139, GLN147, GLN173, GLN177, LEU151, LEU166, LYS169, SER142, SER157, SER160, SER7 |
